# Measuring and Modelling the Emergence of Order in the Mouse Retinocollicular Projection

**DOI:** 10.1101/713628

**Authors:** Daniel Lyngholm, David C. Sterratt, J. J. Johannes Hjorth, David J. Willshaw, Stephen J. Eglen, Ian D. Thompson

**Affiliations:** MRC Centre for Developmental Neurobiology, King’s College London, Guy’s Hospital Campus, London, SE1 1UL, UK; Cyborg Nest Ltd, Clifton House, 46 Clifton Terrace, London, N4 3JP, UK; Institute for Adaptive and Neural Computation, School of Informatics, The University of Edinburgh, 10 Crichton Street, Edinburgh, EH8 9AB, UK; Department of Applied Mathematics and Theoretical Physics, University of Cambridge, Wilberforce Road, Cambridge CB3 0WA, UK; Division of Computational Science and Technology, School of Electrical Engineering and Computer Science, KTH Royal Institute of Technology, Stockholm, Sweden

## Abstract

In the formation of retinotopic maps both experimental and theoretical work implicate guidance molecules and patterned neuronal activity. A common view is that molecular cues *define* and activity cues *refine* mappings. Important insights have come from studies of the retinocollicular projection in transgenic mice, in which cues have been modified either in isolation or in combination. Mostly these have generated descriptions of endpoint mappings. The dynamics of map formation remain under-explored experimentally and computationally. We have quantified changes in the ordering of the mouse retinocollicular projection with age after making local collicular injections of fluorescent microspheres. Contour analysis shows that, at birth (P0), cells from over 80% of the retina converge on a given collicular locus; this percentage falls gradually to P4 then rapidly approaches adult values by P12. Paired injections reveal how the segregation of labelled cells depends both on injection site separation and relative orientation and also age. At P0, large anterior-posterior separations failed to produce segregated label: segregation improved with a similar timecourse to convergence to reach near adult-values by P12. An implementation of a combined activity-molecular model captures these segregation dynamics.

The developmental dynamics were then studied in the nAChR-β2^−/−^ mouse, which has altered patterns of retinal activity in the first postnatal week that results in a more diffuse adult retinal projection. Surprisingly, both measures of map refinement (convergence and segregation) remain largely constant and imprecise in the first postnatal week – only subsequently does the projection begin to refine. Substituting nAChR-β2^−/−^ activity patterns for wild-type patterns [5] into the model failed to capture the biology: refinement was initially *faster*. By reducing the relative importance of the gradients’ contribution to the model energy (to 15% of normal) we were able to mimic the delayed refinement observed in vivo. Our model therefore predicts the altered activity patterns may affect readout of guidance cues.

## Introduction

Topographically precise neural maps are a widespread feature of the nervous system that facilitate neural networks to convey positional information across different neural systems for parallel processing. In particular, such topographical organisation is found throughout the visual system, where retinal neighbour relationships are preserved in primary and secondary visual structures. The relative contribution of molecular cues and instructive spontaneous activity during map formation, and whether they are independent processes or somehow interact, is largely unknown. Further, it is unclear that our current understanding of actual molecular components and spontaneous or sensory modulated activity represents the full complement of mechanisms involved at all scales of the constructed circuit (Cang and Feldheim, 2013; Arroyo and Feller, 2016; Ito and Feldheim, 2018).

Although numerous studies have provided an insight into the development of the retinocollicular projection (Hindges et al., 2002; McLaughlin et al., 2003; Mrsic-Flogel et al., 2005; Rashid et al., 2005; Cang et al., 2008; Xu et al., 2011, 2015; Burbridge et al., 2014; Owens et al., 2015; Savier et al., 2017), these studies largely focused on a restricted period of development, or were qualitative. There is a lack of quantitative data on the spatiotemporal sequence of development, which is needed for making predictions about how patterns of guidance cues and activity patterns constrain development of spatially precise topographic projections.

To examine the role of patterned activity in the early development of topography we have utilised a global knockout of the β2 subunit of the nicotinic acetylcholine receptor (nAChR-β2). This mutant has altered spontaneous activity patterns in the retina and the β2-subunit is therefore believed to be crucial for generating the specific pattern of correlated activity waves seen in the retina from P0-P8 (Bansal et al., 2000; Sun et al., 2008) that propagate throughout the visual system (Ackman et al., 2012). Moreover, this mutant has mapping defects in the visual system (McLaughlin et al., 2003; Grubb and Thompson, 2004; Mrsic-Flogel et al., 2005; Rashid et al., 2005; Cang et al., 2008; Shah and Crair, 2008; Burbridge et al., 2014). Importantly, the spatiotemporal characteristics of the waves in both wild type and nAChR-β2^−/−^ are well-characterised (Sun et al., 2008; Stafford et al., 2009; Xu et al., 2011) and can therefore be used in models to make predictions about the precise patterns of activity needed for the instructive roles of the waves.

Here we provide an in-depth quantitative spatiotemporal description of the postnatal development of topographical organisation in the mouse retinocollicular projection by comparing point-to-point connectivity between the retina and superior colliculus (SC) using a novel approach for reconstructing and standardising retinal flat-mounts (Bansal et al., 2000; Sun et al., 2008; Sterratt et al., 2013). For the antero-posterior (AP) axis of the SC, we show that only limited topographical precision is present at P0 and that precise topography only starts to appear after P4 and reaches adult levels by around P12. Moreover, we demonstrate how the topographic refinement between P4-P8 is dependent on specific spatiotemporal retinal activity patterns, as altering these prevents establishment of precision in the first postnatal week and results in a marked decrease in topographic precision, which is never fully recovered in adulthood. Finally we demonstrate by using a modelling approach (Triplett et al., 2011; Hjorth et al., 2015) that the interactions between guidance cues and activity are crucial for establishing precision.

## Results

The adult mammalian retinocollicular projection is characterised by a high order of topography, where adjacent retinal ganglion cells (RGCs) project to adjacent areas in the SC.

### Quantification of precision in mature and immature projection

In the mature visual system, as can be seen in Figure 1A, two calibrated (5-10 nl) injections separated by 65 µm along the AP axis of the SC can give two distinct foci of labelled RGCs with only limited overlap. In contrast to this, at birth, paired injections separated by 1200 µm along the SC AP axis result in almost complete overlap of the two tracers in the retina (Figure 1B). However, while most of the retina contains labelled cells, there is still a level of AP order in the projection at birth. Figure 1B shows that the Karcher mean retinal location of cells labelled from the more anterior injection (blue diamond) is located in temporal retina whereas that for the more posterior injection (red diamond) is located in nasal retina. The retinal vectors linking the centres of retinal label (either Karcher mean or Peak density) for paired antero-posterior injections in P0 animals (n = 11) were all oriented appropriately. It is known from studies in many animals including the rodent (Simon and O’Leary, 1992; Plas et al., 2005) that there is topographic order for the dorsoventral (DV) retinal axis in the optic tract. Thus, unsurprisingly, the distributions of labelled RGCs resulting from injections separated by 1070 µm along the SC medio-lateral (ML) axis are segregated (Figure 1C). In the mature visual system a single discrete injection of tracer into the superficial SC results in a defined focus of label that covers 3.6±2.2% (all values are given as mean ± standard deviation) of the retina surface (Figure 1D). In contrast to this, the projection at birth is very immature and therefore a similar discrete injection will result in 76.9±9.5% of the retina containing labelled cells (Figure 1D). The most densely packed labelled RGCs occupy much less of the retina in both adult and neonatal examples (75% contour – adult: 0.23±0.15% and P0: 10.2±2.6%). The use of paired injections allows investigation of the precision of the projection at multiple ages and across the two SC axes. When quantifying the segregation of retinal label (see Methods) and plotting this as a function of injection site separation, the resulting curves reveal two marked differences between the adult and P0 data. The adult curves rise very rapidly to give complete segregation at relatively small injection separations and shown no axial differences. The P0 curves are much flatter and display marked axial differences. There is an obvious difference given the non-overlapping confidence intervals of the functions fitted to the ML and AP populations (Figure 1E), indicating considerable anisotropy in the projection. The absence of such a difference in the mature projection indicates that it is isotropic.

**Figure 1:**
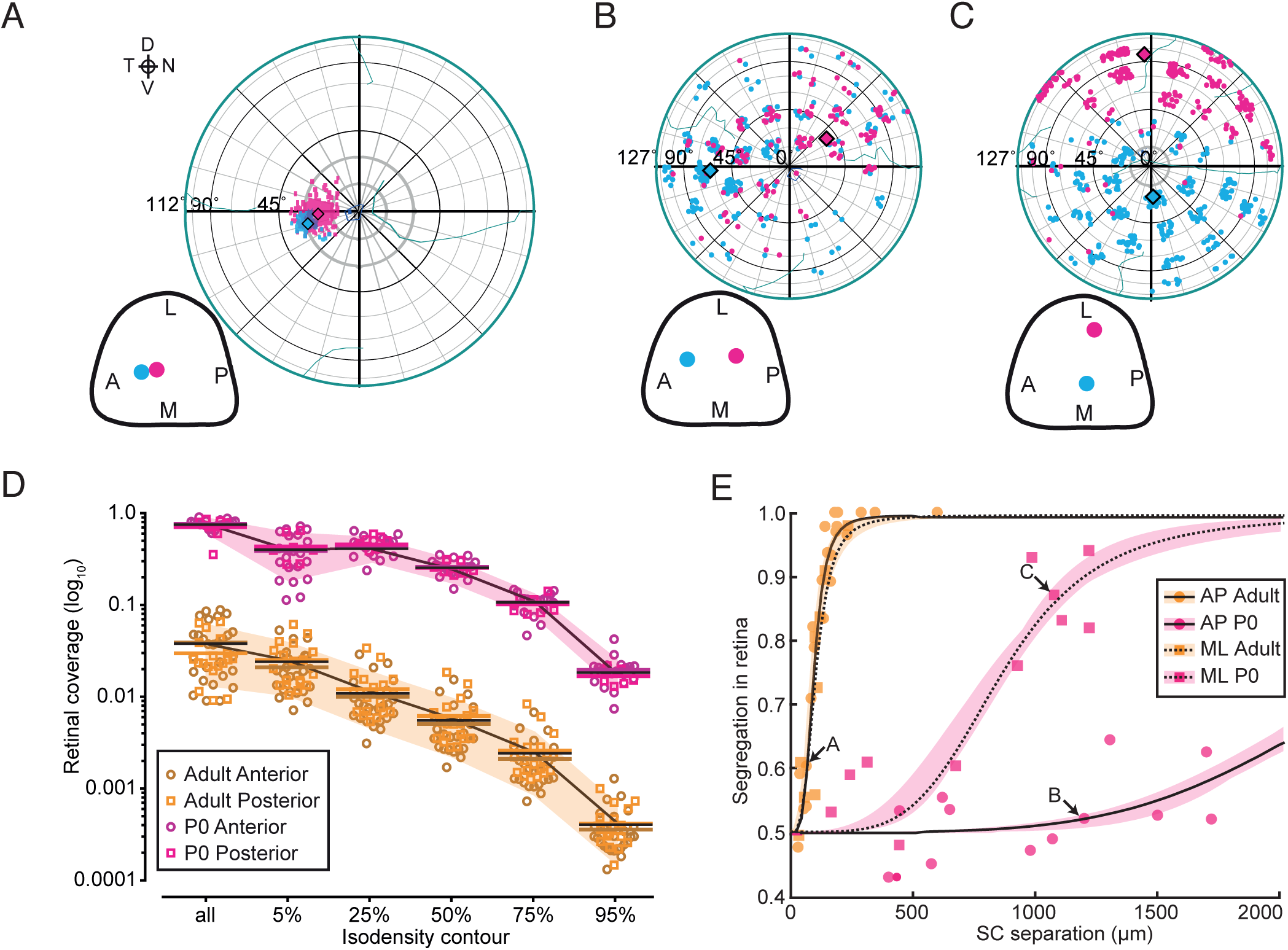
Adult and neonatal precision. **A-C**, Examples of retinal label from paired injections into the adult SC separated along AP SC axis (A) and into the neonatal SC separated along AP (B) and ML (C) SC axis. Retinae have been reconstructed and projected in an azimuthal equidistant projection (Sterratt et al., 2013). Insert shows the locations of the injection sites in the SC for each retina. **D**, Amount of retina containing labelled cells (all) and area covered by isodensity contours for 5%, 25%, 50%, 75% & 95% of the peak density of retinal label for Adult (magenta) and P0 (orange). Coverage values are shown independently for injections into anterior SC (circles) and posterior SC (squares). The means are represented by coloured lines in corresponding shades. Black lines indicate the mean for respective age and shaded area is the standard deviation. **E**, Nearest-neighbour value (see Methods) for paired injections plotted as a function of SC injection site separation along AP (circles) and ML (squares) axes. Curves for AP (solid lines) and ML (dashed lines) axes are fitted with a Hill function using a jackknife approach as described in Methods. Shaded area represents the distribution of individual jackknife fits. Arrows indicate the values for the examples shown in A-C.

### Developmental dynamics of projection precision and order

Because of the significant refinement of the projection, especially along the AP axis, a key question is what are the precise spatiotemporal dynamics of this refinement. Figure 2A shows representative examples from ages sampled during the first 3 postnatal weeks with isodensity contours plotted for cells at 25%, 50% and 75% of the peak density. At P2, the label resulting from a single discrete injection still covers the entire nasotemporal (NT) axis but, just like at P0 (Figure 1B-C), it is more defined in the DV axis. At P4, this anisotropy is largely gone and the label now covers equal proportions of the DV and NT axes. At this age a focus is, moreover, visible. By P6, despite a large spread of peripheral label, a clearly defined symmetric focus is evident and as a result the peak density and Karcher mean of the label coincide. The peripheral labelled cells persist at P8, but are gone by P12. The quantification of isodensity contours is shown in Figure 2B. The relative extent of retina containing any labelled cells changes very little over the first four postnatal days with a steep fall between P4 and P6. In contrast, the focus of label as defined by the 25% isodensity contour shows a different pattern. There is a significant (*p*<0.0001; one-way ANOVA with Bonferroni test of multiple-comparisons) decrease between P0 and P2. The curves for 25%, 50% and 75% isodensity contours show a rapid refinement to P6 with smaller subsequent changes. The data for all cells remains relatively expanded up to at least P12.

**Figure 2:**
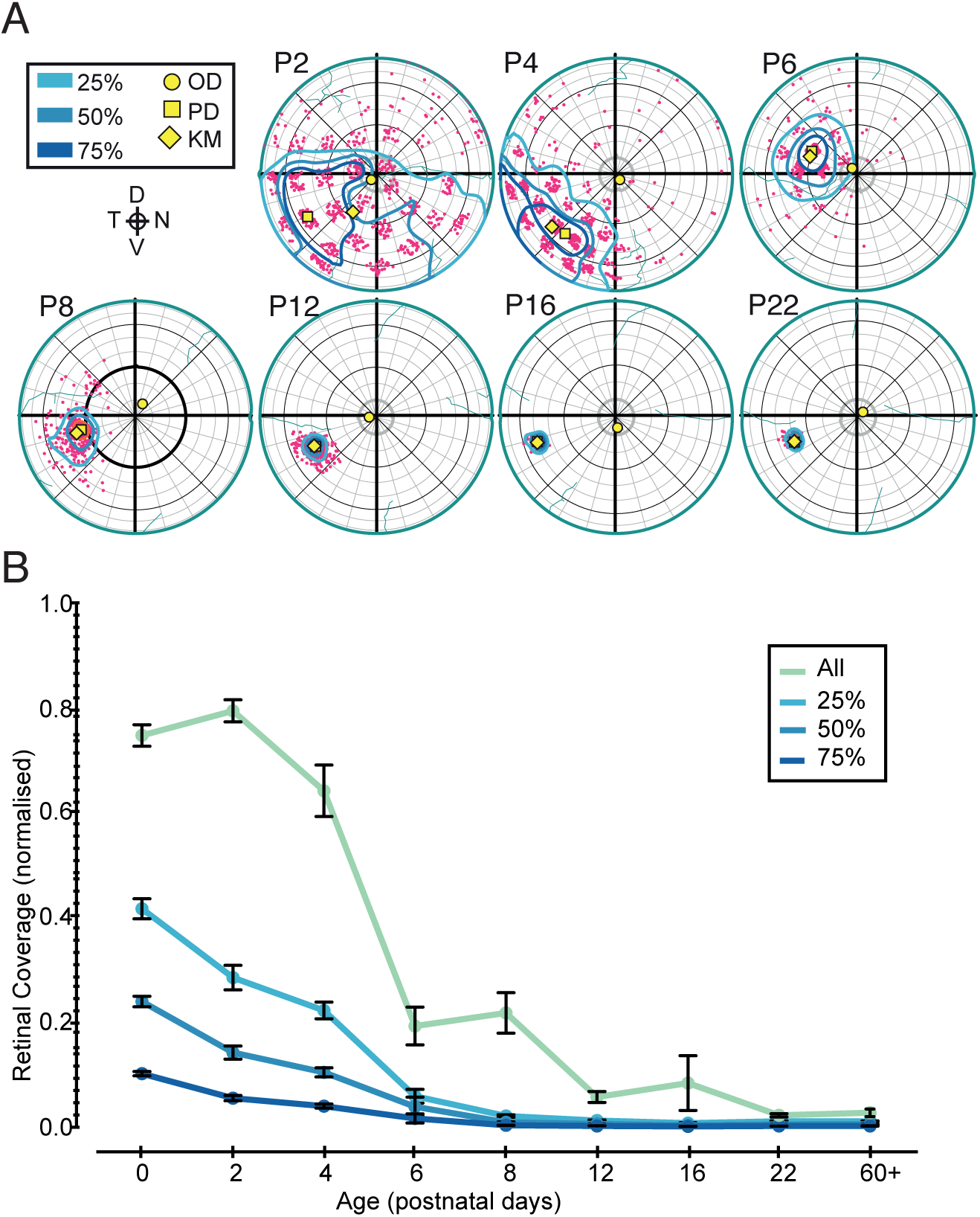
Development of precision. **A**, Examples of label from single injections into the SC at given ages. Examples for P2, P4 & P6 were partially sampled (see Methods) and P8+ were sampled completely. Isodensity contours were plotted for 25%, 50% and 75% of peak density. The Karcher mean (KM) is indicated with a diamond, the peak density (PD) is indicated with a square and the optic disc (OD) is indicated with a circle. Retinal orientations as in Figure 1. **B**, The mean proportion of retina containing labelled cells (All) and the area covered by the isodensity contours in A at given postnatal ages. Error bars are SEM.

The isodensity contour analysis in Figure 2 provides a good description of the convergence of retina to SC. Figure 3 shows another measure of mapping precision: the segregation in retina derived from paired injections at different separations on the SC. When quantifying the segregation of retinal label (see Methods) for paired injections along the AP axis of the SC (Figure 3A) there are significant differences in the function fits between contiguous ages from P0 to P12 with the exception of P6 to P8, which are very similar. Surprisingly, the functions fitted for P22 and adult (60+) are also different. For the injections paired along the ML axis of the SC (Figure 3B) the starting point is different to the AP axis, since DV RGC axons are ordered in the brachium of the SC (Plas et al., 2005). Despite differences in the starting precision, the developmental dynamics of refinement along the ML axis appear similar to that for AP axis: the biggest shift in the curves is between P2 and P4 and between P4 and P6/P8. By P12, the curves in Figures 3A and 3B are very similar: the early anisotropy (Figure 1B and C) has been lost. For the ML axis there is also a difference between the fitted functions for P22 and adult (60+). Figure 3C is another measure of the developmental dynamics. From the fitted curves in Figures 3A,B can be derived the SC separation that corresponds to a 95% segregation index (dotted lines in Figures 3A,B). There are clear differences in the absolute values at the earliest ages, but the dynamics for this measure show parallel changes for both axes between P4 and P8. There is further refinement along the AP axis between P8 and P12, after which the projection is isotropic.

**Figure 3:**
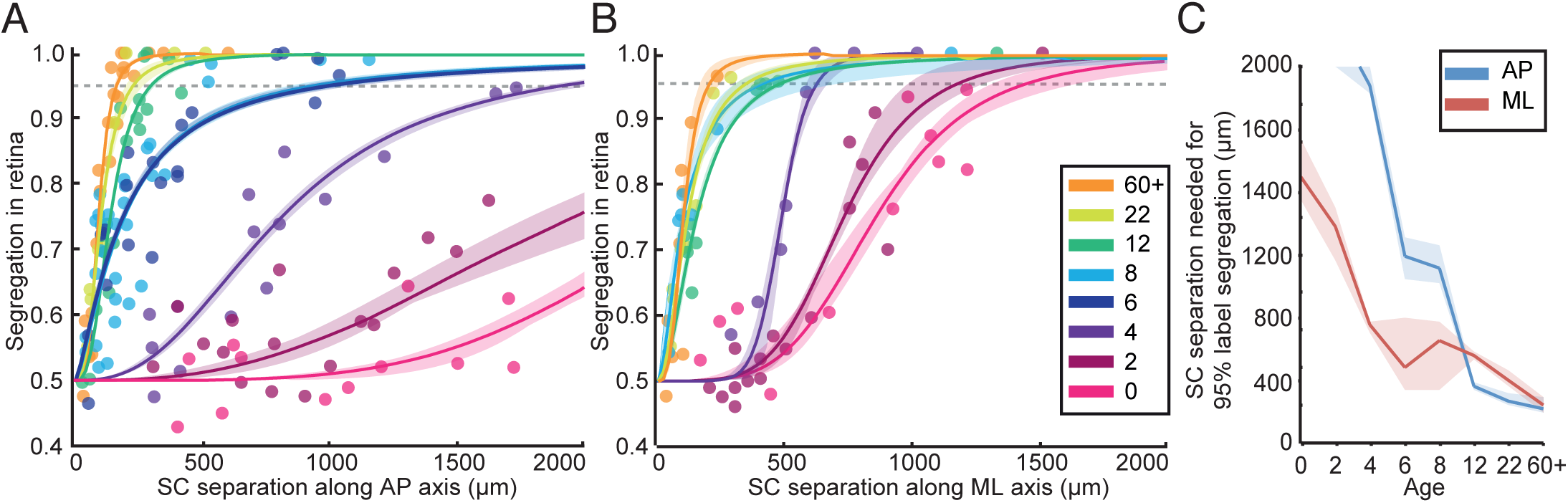
Development of order. **A-B**, Hill function fits (as in Figure 1E) to label segregation analysis for paired SC injections separated along the AP axis (A) or ML axis (B) for given developmental ages and adult (60+). Shaded area represents the accuracy of the fits. Note that the fitted curves for the P6 and P8 data substantially overlap in both A and B. Dotted lines indicate 95% segregation in retina. **C**, The corresponding SC separations for different postnatal ages.

### Modelling development of the normal projection

A retinotopic map formation model (Triplett et al., 2011; Hjorth et al., 2015) was implemented to distinguish between two hypotheses: that a change in retinal activity in nAChR-β2^−/−^ mice impairs Hebbian style refinement of connections, and that altered activity inhibits readout of chemical cues. The development of the wild type was modelled, using parameters for retinal activity described for wild type in Stafford et al. (2009) (Figure 4A, see Methods). Virtual paired injections were made with various separations, and the data used to create segregation curves. This procedure was repeated at various iterations of the model to generate groups of segregation curves. Figure 4B shows how the iteration curves match the real data curves for AP injections (Figure 3A). For any age, there is an iteration when the match between model and real data is minimal. The arrows in Figure 4B show the iteration number giving the best match at a given age. Figure 4C shows virtual tracer injections at two locations in the model SC, and the corresponding model distributions across the retina for the matched iterations are shown in Figure 4B. Qualitatively, the modelled projection gradually refines as in wild type (Figure 2A). Figure 4D illustrates how the modelled segregation curves (solid lines) match the real data. The model can capture the developmental dynamics of the actual data when matched to the retinal segregation index. But does it also capture the changes in coverage derived from isodensity contours? The overall shapes of the segregation fits are similar, but there are differences between experimental data and model. A potential problem for the age to iteration matching approach is that the experimental P6 and P8 curves overlap, and will match to the same or very close iterations in the model. This is shown in Figure 4E. In both model and data, the majority of the refinement, based on convergence measures occurs before P6. Note that, before P6, the modelled convergence values (solid line) are higher than the data values (dotted line). This is because there is no early anisotropy in the model projection, unlike the *in viv*o state.

**Figure 4:**
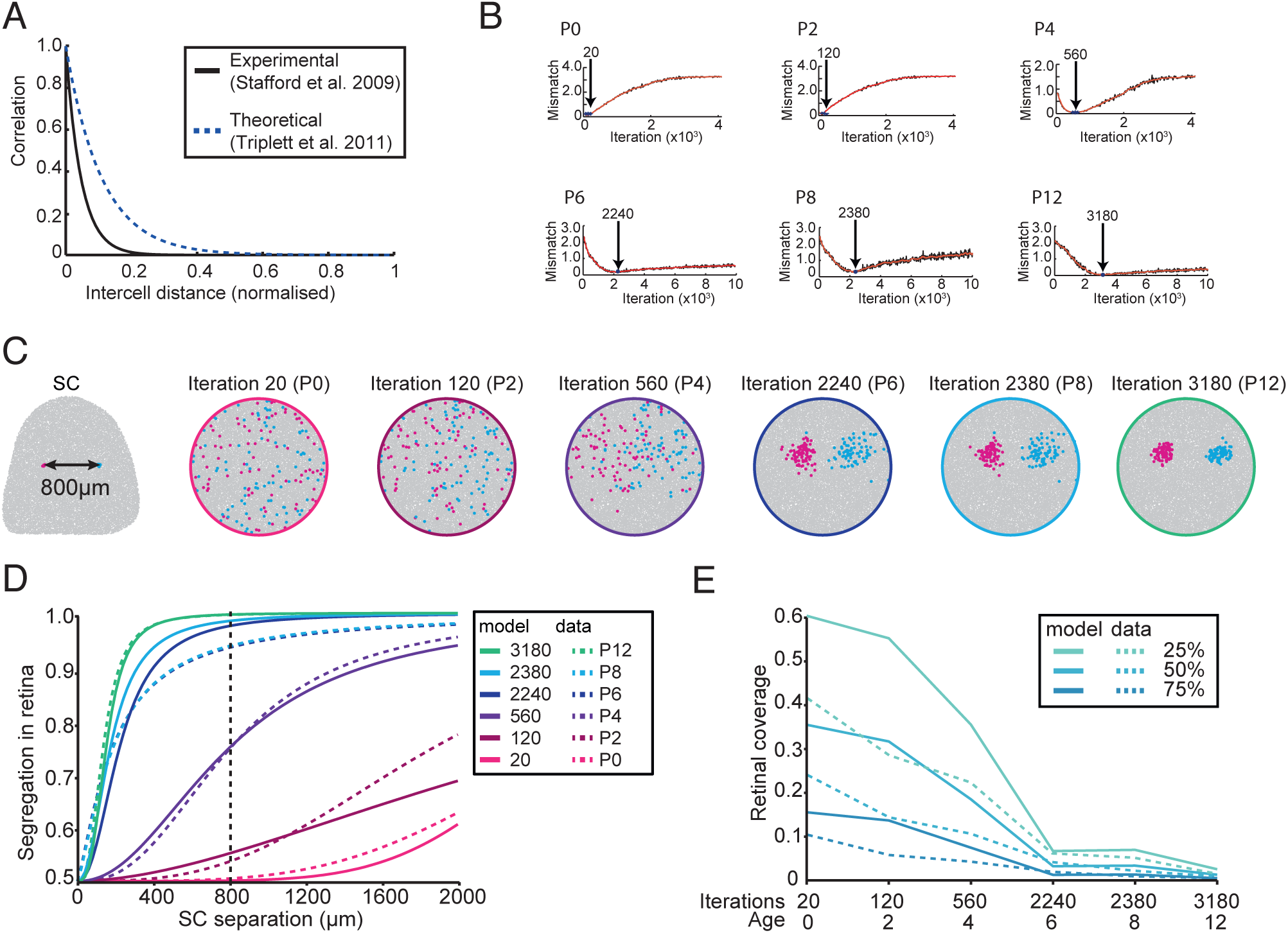
Modelling Development. **A**, Experimental correlation profiles from Stafford *et al.* (2009) used in this model (solid black line) and theoretical correlation profile used in the implementation of model by Triplett *et al*. (2011). **B**, Matching of iteration of model to experimental age for P0 to P12. Plot shows is of mismatch of simulated segregation curves to the data of the given age, where 0 is a perfect fit. Arrows indicate the minimum mismatch. **C**, Examples of simulated paired retrograde injections into the SC separated at 800 µm along the AP axis at the iterations equivalent to ages in B. **D**, Segregation in retina functions for simulated and matched experimental data for P0 to P12 (dashed lines). Vertical dashed line indicates separation of virtual injections in C**. E**, Retinal coverage associated with simulations shown in B-C (solid lines) plotted with equivalent matched experimental data for ages P0 to P12 (dashed lines).

### Comparing mature and neonatal nAChR-β2^−/−^ animals

To assess the influence of correlated spontaneous activity in establishing a topographically precise projection, we have performed the experiments and analyses described in Figures 1-3 for the nAChR-β2^−/−^ mouse line. These animals have retinal spontaneous activity but with altered spatiotemporal correlations (Stafford et al., 2009). When examining the adult, it is evident that the projection is noticeably less precise: the foci observed in Figure 5A are much larger than those in Figure 1A. Moreover, although the injection separation (192µm) in this animal was much greater than the animal shown in Figure 1A (65µm), there is almost total overlap of the two foci of label. For animals injected at P0 (Figure 5B), on the other hand, the projection appears similar to wild type (Figure 1B). The isodensity analysis shows that in the nAChR-β2^−/−^ adult, a single discrete injection of tracer into the SC results in a focus of label that covers 18.3±10.5% of the retinal surface (compared to 3.6±2.2% in wild type) whereas a similar injection in nAChR-β2^−/−^ P0 labels 82.5±11.7% of the retina (compared 76.9±9.5% in wild type). The most densely packed labelled RGCs occupy much less of the retina in both adult and neonatal examples (75% contour: 1.8±1.0% adult and 7.3±2.5% P0). Although the isodensity contours for P0 and adult are significantly different (*p*<0.0001), the mean difference is less than for wild type (Figure 5C). When investigating the label segregation in nAChR-β2^−/−^ animals (Figure 5D), we find that there is no real difference between the fitted functions at P0, whereas there is a pronounced difference in adult animals.

**Figure 5:**
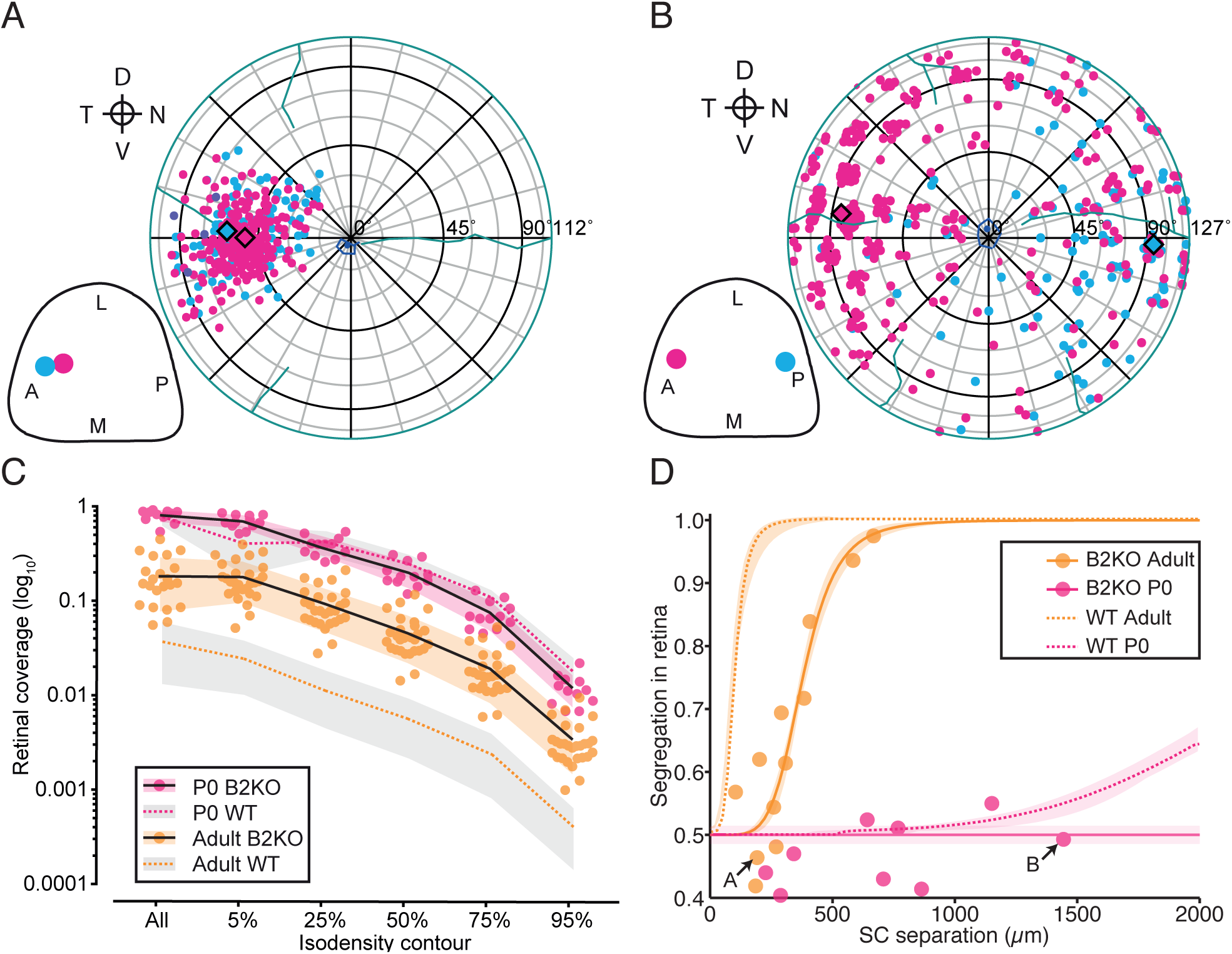
Adult and neonatal precision in nAChR-β2^−/−^ animals. **A-B**, Example of retinal label from paired injections into the adult SC (A) and neonatal SC (B) separated along AP axis. Insert shows the locations of the injection sites in the SC for each retina. **C**, Amount of retina containing labelled cells (All) and area covered by isodensity contours for 5%, 25%, 50%, 75% & 95% of the peak density of retinal label for Adult (orange) and P0 (magenta). Black lines indicate the mean for respective age and Coloured dashed lines represent the corresponding data for wild-type animals from Figure 1D. Shaded area is the standard deviation. **D**, Segregation in retina for paired injections plotted as a function of SC injection site separation along AP axis. Fitting of functions as in Figure 1E. Arrows indicate the values for the examples in A-B. Dashed lines represent the corresponding fits for wild-type animals from Figure 1E.

### Developmental dynamics of projection precision and order in nAChR-β2^−/−^ animals

The developmental dynamics of map refinement in the nAChR-β2^−/−^ mice are illustrated in Figure 6. Between P2 and P6 labelled cells still span most of the retinal NT axis and, although 50% and 75% isodensity contour lines do show some hemiretinal restriction at P6, there is no evidence for the rapid refinement of mapping along the AP axis of the SC that was observed in wild type (see Figure 2). In the examples illustrated there is a big difference between P8 and P12. Quantifying the above isodensity contours (Figure 6B) reveals that, in contrast to wild type, the slope of the contour plots changes very little between P0 and P6. There are no noticeable decreases in the retinal coverage in adjacent ages up to P6. From P6 to P8, there is a gradual refinement in coverage, although the overall spread of labelled cells (‘All’) remains elevated. The difference could imply a different dynamic for refinement of focal and ectopic (as revealed in the ‘All’ data cells). Figures 6C and 6D compare retinal coverage in the nAChR-β2^−/−^ and wild type across ages and contour lines. There is little difference in the coverage ratios at P0 (Figure 6C). The gradual subsequent rise at P2 and P4 in the coverage ratios for 25%, 50% and 75% isodensity contours reflects refinement in wild type and stasis in the mutant (compare Figures 2B and 6B). Elimination of ectopic cells begins after P2 in wild type, explaining the ‘All’ profile in Figure 6C. Later phases of development are shown in Figure 6D. The peak in coverage ratio at P8 for the 25%, 50% and 75% contours reflects the drop in retinal coverage to near adult values at this age in wild type and the absence of any difference between P6 and P8 for coverage in the mutant. The removal of the ectopic cells (‘All’ contours, Figure 6D) reflects (i) relatively delayed removal of such cells in wild type and (ii) their relative persistence in the nAChR-β2^−/−^ (compare Figures 2B and 6B).

**Figure 6:**
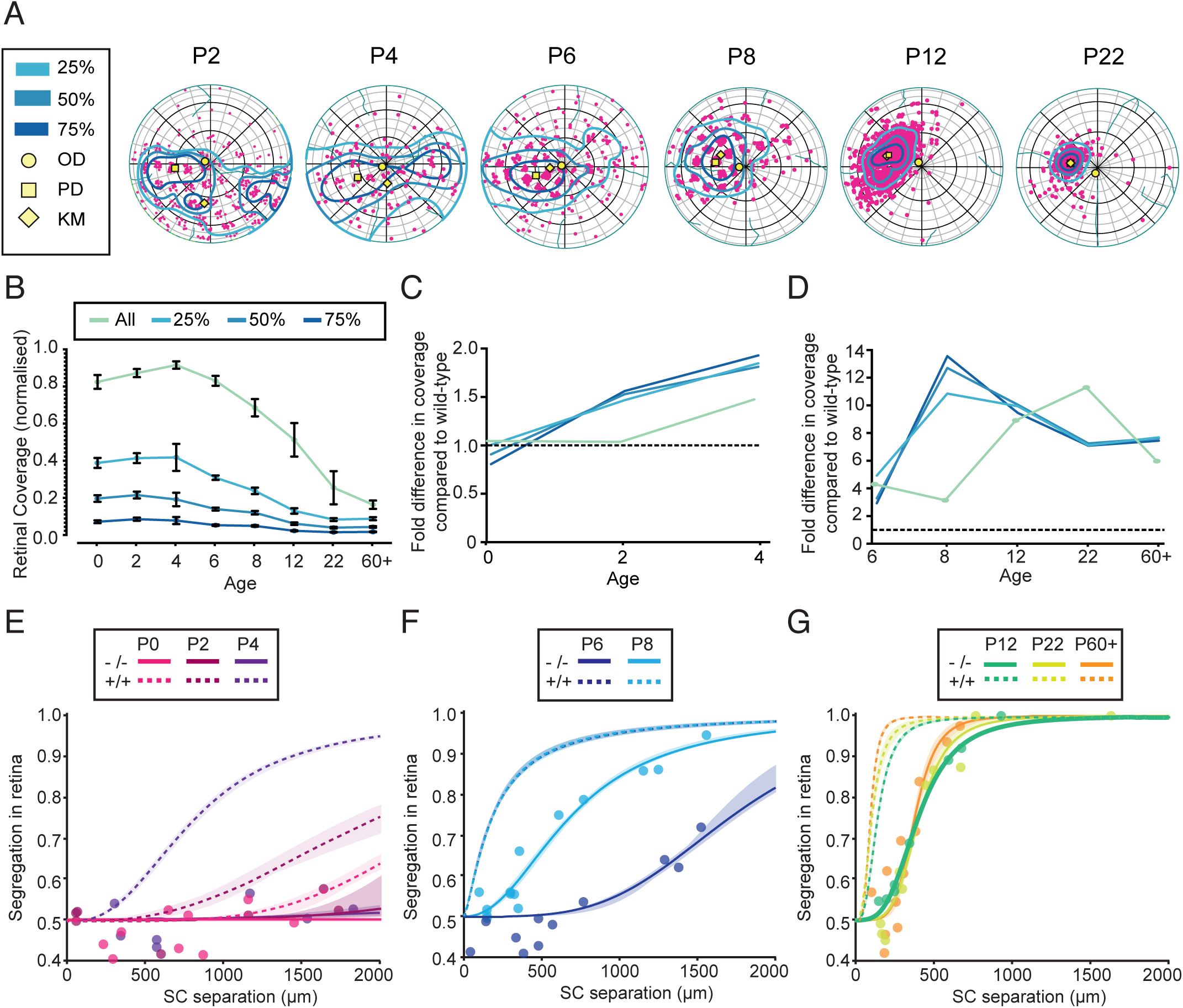
Impaired development of precision and order in nAChR-β2^−/−^ animals. **A**, Examples of label from single injections into the SC at given ages. Examples for P2, P4, P6 and P8 were partially sampled whereas P12, P22 and adult were sampled completely (see Methods). Isodensity contours were plotted to 25%, 50% & 75% of peak density. The Karcher mean (KM) is indicated with a diamond, the peak density (PD) is indicated with a square and the optic disc (OD) is indicated with a circle. Retinal orientations as in Figure 5A **B**, The mean proportion of retina containing labelled cells (All) and area covered by the isodensity contours in A. Error bars are SEM. **C**, Fold increase in retinal coverage of nAChR-β2^−/−^ animals, compared to wild type for P0-P4. **D**, Fold increase in retinal coverage of nAChR-β2^−/−^ animals, compared to wild type for P6+. Dashed lines in C and D indicate equivalence between wild-type and mutant coverage. **E-G**, Hill function fits to segregation in retina for paired SC injections separated along the AP axis for P0-4 (E), P6-8 (F) and P12-60+ (G). Shaded areas reflect the uncertainty associated with the jackknife fits (see Figure 1E). Solid lines are nAChR-β2^−/−^ and dashed lines are wild type.

For paired injections separated along the SC AP axis, the segregation in retina distributions (Figure 6E) shows that in the nAChR-β2^−/−^ animals there is no difference between the functions fitted at P0, P2 or P4. Despite there being no significant difference between the isodensity contour coverages for P4 and P6 animals, the segregation plots indicate that there is some refinement in the AP axis at P6, which becomes more marked by P8 (Figure 6F). In Figures 6E-G, the segregation plots for wild type animals are given by the dotted lines. Comparing curves, the fit for P6 nAChR-β2^−/−^ is most similar to P2 wild type and the fit for P8 nAChR-β2^−/−^ closely resembles that for P4 wild type. In contrast to wild type, there is a large difference between P6 and P8 curves in the nAChR-β2^−/−^ animals, whereas the pattern of fits for P12 to P60 has a very similar shape to wild type, albeit shifted significantly by 250µm.

### Modelling development of the nAChR-β2^−/−^ projection: elucidating the contributions of correlated activity

The nAChR-β2^−/−^ simulations differ from wild type only in the correlation function used, with a wider spatial correlation and a lower peak (see Figure 7A). Using the iterations that correspond to the ages P0-P12 in wild type, the maps at corresponding ages for the nAChR-β2^−/−^ model can be investigated. Figure 7B shows simulated retinal label distributions in the mutant following virtual injections in the SC at 800 µm separation The simulated map refines continuously and appears very similar to the wild-type simulation (Figure 4C) that matches the wild-type developmental data. The simulation is very different from the developmental data from the nAChR-β2^−/−^ at the early ages, which shows a clear delay in refinement (Figure 6A). In fact, plots of the simulated retinal segregation (Figure 7C) are shifted to the left, implying an *accelerated* development of order (compare Figure 7C and Figure 4C). To emphasise this point, Figure 7D illustrates the simulated segregation values for an injection separation of 800 µm for wild type and mutant. The segregation values are consistently higher in the mutant, especially at iterations equivalent, in wild type, to P4. Figure 7E compares simulated isodensity retinal coverage: again, at P4, the mutant simulation shows a more focussed projection retinal coverage than the wild-type simulation.

**Figure 7:**
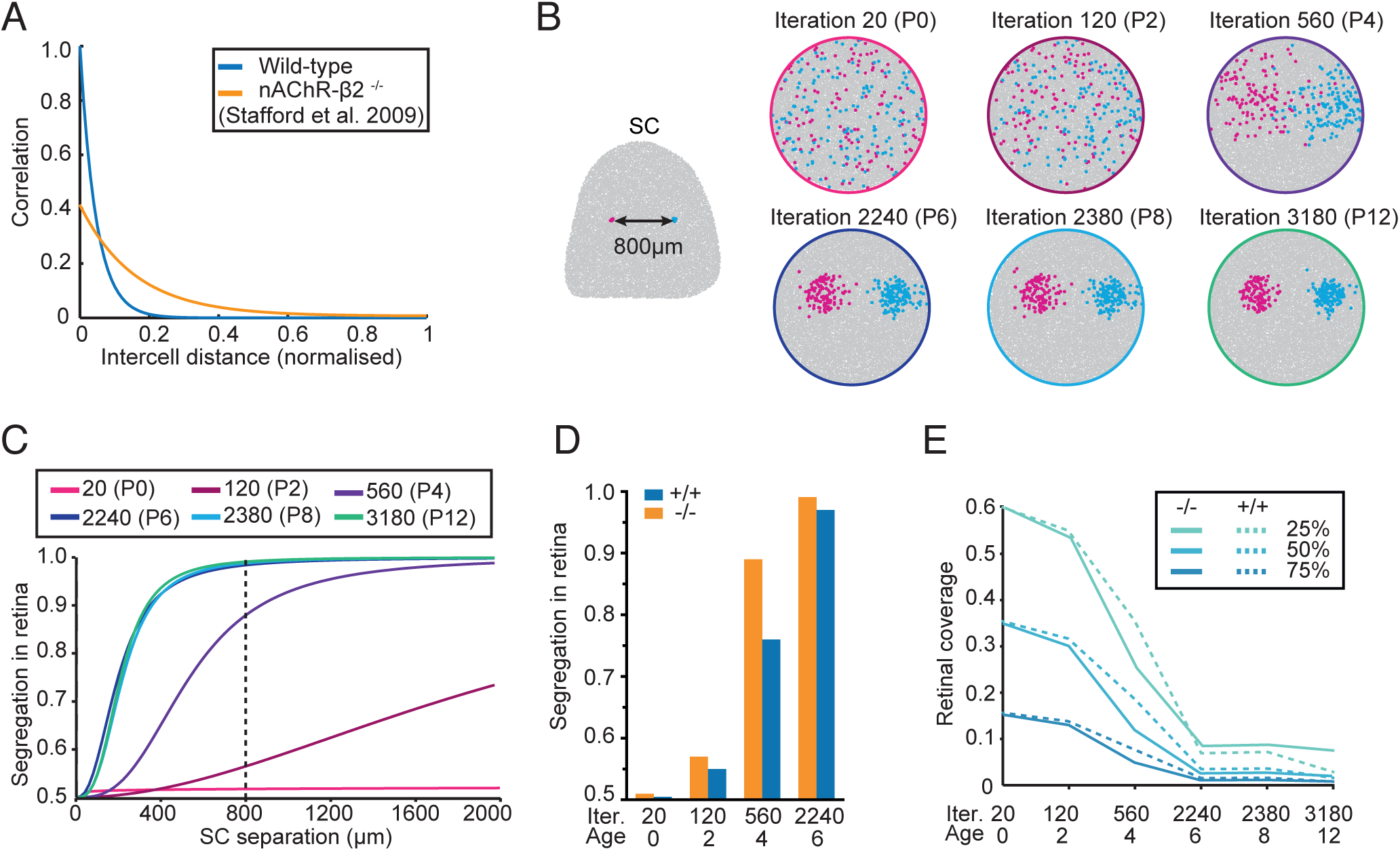
Modelling the dynamics of nAChR-β2^−/−^ development. **A**, Correlations used in the model. Data from Stafford *et al.* (2009) for wild-type and nAChR-β2 ^−/−^ retinal in vitro explants. **B**, Example of simulated paired retrograde injections through development (using nAChR-β2 ^−/−^ correlations in A) at iterations equivalent to ages from P0 to P12 using the wild type age-to-iteration matching (in Figure 4B). Virtual injection separation was 800µm. **C**, Segregation in retina for simulated nAChR-β2^−/−^ animals. Vertical dashed line indicates separation of virtual injections (800 µm) in B. **D**, Simulated segregation in retina for iterations equivalent to wild-type P0,2,4,6 and an injection separation of 800µm. Orange indicates nAChR-β2 ^−/−^ simulations and blue wild-type simulations. **E**, Retinal coverage for simulated nAChR-β2^−/−^ at iterations in B (solid line) compared to equivalent coverage from wild-type simulations (from Figure 4E; dashed line) at matched iterations.

### Modelling development of the nAChR-β2^−/−^ projection: elucidating the relative contributions of molecular cues

Our earlier modelling results suggested that the Triplett et al (2011) model accounted for retinotopic maps under a range of experimental conditions (Hjorth et al., 2015). However, this model could not explain the dynamics of refinement of the nAChR-β2^−/−^ map by suitable adjustment of retinal correlations in the model. One hypothesis is that the change in activity also acts indirectly by reducing the readout of the gradient information (Nicol et al., 2007). To model this, the readout of the chemical cues was decreased by introducing a scale factor in the molecular component of the model. We chose to look at simulations of the segregation in retina after 560 iterations, corresponding to wild-type P4: at this stage the differences between wild type and mutant simulations and wild type and mutation data are greatest (Figure 8A). Figure 8B illustrates how, using the spatiotemporal correlations for the nAChR-β2^−/−^, decreased chemical readout affects the segregation in retina values. Not surprisingly, as the chemical cue is weakened the emergence of topography is also weakened. By 15% of the original chemical cue strength the plot of virtual segregation versus virtual injection separation at iteration 560 (=P4 wild type) is comparable to actual P4 nAChR-β2^−/−^ data: i.e. almost no segregation. One clear result was that reducing the strength of chemical cues meant that the simulations took longer to converge to mature maps (Figure 8C).

**Figure 8:**
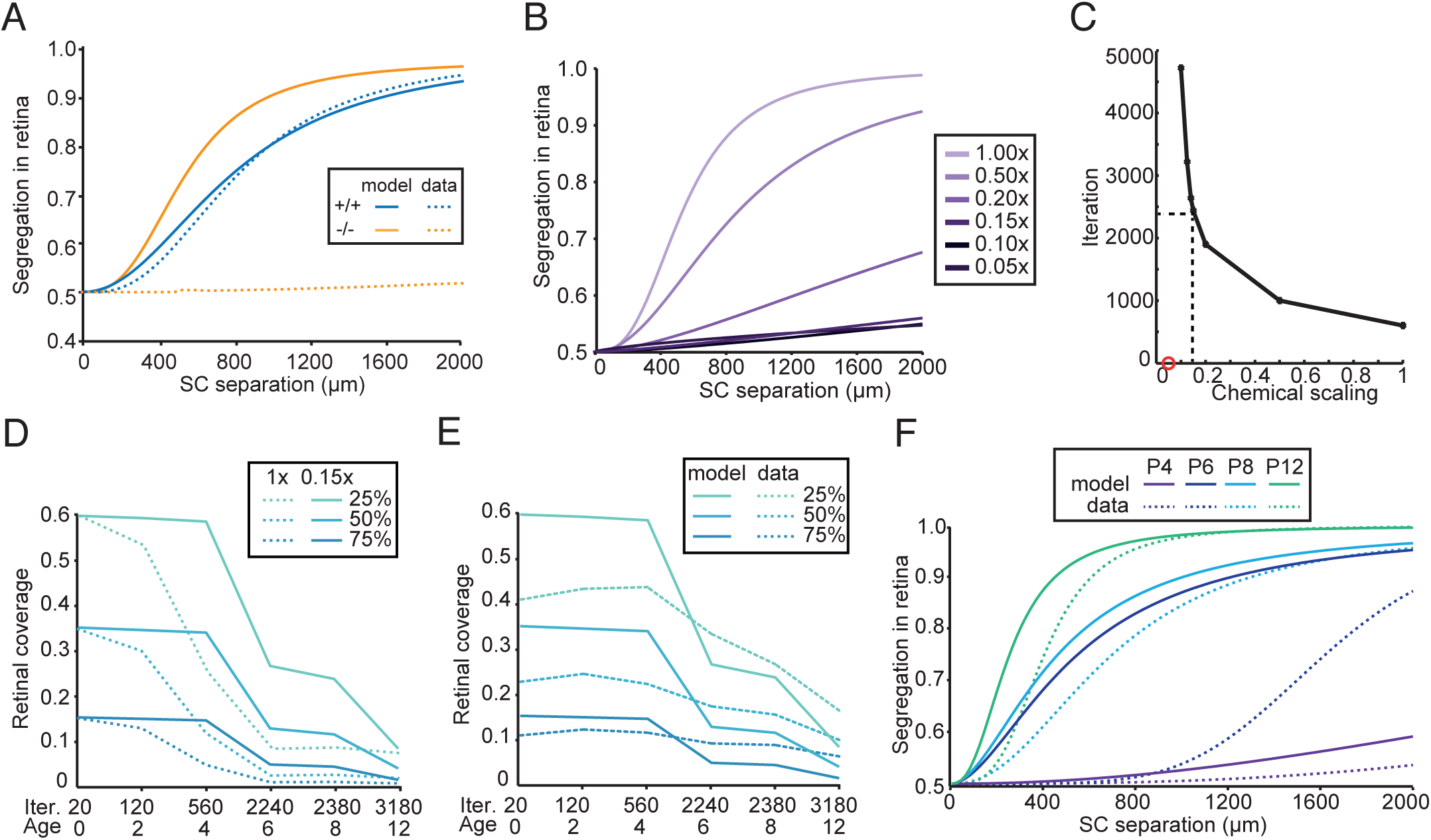
Elucidating the relative contribution of molecular cues. **A**, Segregation in retina as a function of distance in the SC of simulated data at 560 iterations and equivalent experimental P4 data for nAChR-β2^−/−^ and wild type. **B**, Simulated segregation in retina for various scalings of molecular cues at 560 iterations. **C**, Iterations needed for a simulation with reduced chemical cues to reach the same segregation precision as a default simulation. Red circle indicates that this did not happen within 10,000 iterations. Dashed line indicates a scaling of 0.15x, which is the best fit to P4 data in A. **D**, Retinal coverage for simulated nAChR-β2^−/−^ animals with molecular cues scaled at 0.15x (solid lines) compared to iteration-matched values for molecular cues at 1.0x (Figure 7D). **E**, Comparison of retinal coverage values of simulated nAChR-β2^−/−^ animals scaled at 0.15x (solid lines) with age-matched nAChR-β2^−/−^ anatomical data. **F**, Comparison of segregation in retina for simulated nAChR-β2^−/−^ animals scaled at 0.15x (solid lines) with age-matched nAChR-β2^−/−^ anatomical data.

Figure 8D compares simulations of the dynamics of retinal convergence with chemical cues at full-strength and at 15%. Weakening the chemical cues does delay map precision, with big differences at iterations equivalent to P4 and P6; the curves converge by the equivalent of P12. The final two panels (Figures 8E,F) compare the convergence and segregation indices for the new simulation and the nAChR-β2^−/−^ data. For the retinal segregation index, the agreement between the data and the new model is much better than with the old model. The exception is the P6 data. This arises because the equivalent iterations for P6 and P8 were derived from the wild-type data, which showed little difference in segregation values at the two ages, generating very similar iteration values. As with Figure 4E, the convergence values cannot be compared on the basis of absolute values, but the new simulated curves fit the real data much better than the old model. Again there is a problem with the P6 data as above.

Figure 8E shows the development of the nAChR-β2^−/−^ model using 15% of the chemical strength. The refinement of the map initially stalls and iteration 560 matches P4 nAChR-β2^−/−^ data. By iteration 2380 it matches the more refined nAChR-β2^−/−^ P8 data, and the precision at iteration 3180 is comparable to nAChR-β2^−/−^ P12. The only discrepancy is the matching of the model to nAChR-β2^−/−^ P6, which stems from how we match age to iterations and because wild-type P6 and P8 are similar, giving them very close corresponding iterations (2240 and 2380). We conclude that the delay can be accounted for in the model by weakening the readout of chemical cues.

## Discussion

We have provided a quantitative description of the development of anatomical precision in the retinocollicular projection showing that there are multiple stages to wild-type development, which differ significantly for the cardinal axes of the SC. Moreover, we show that altered spontaneous retinal activity with a wider correlation index (Stafford et al., 2009) results in a delayed development of precision after the first postnatal week that is not recovered in adulthood, suggesting a critical role for specific retinal spontaneous activity in early postnatal development. Next we demonstrate that using wild-type correlation indices from Stafford et al. (2009) to constrain a model of topographic development (Triplett et al., 2011) results in a projection comparable to the anatomical data. Using this model we suggest the possibility that the altered projection resulting from the broader activity-correlation index could be due to an impaired read-out of molecular cues.

The quantitative approach adopted here reveals several unexpected features of the topographic development of the wild-type retinocollicular projection. While the difference in precision for the AP versus ML axes of the SC was not unexpected, given previous DiI studies (Hindges et al., 2002), data from single injections show that RGCs from across the entire retina converge on a single SC location at P0, although the distribution is not uniform. Data from injections paired across the AP axis of the SC at P0 generate two, apparently contradictory, results: (i) the relative locations of the *peaks* of labelling in the retina have an appropriate nasotemporal polarity, suggesting a crude map; and (ii) that the segregation of labelling in the retina is independent of injection separation, suggesting no map. Note that values for segregation in the retina rely on the *total* population of labelled RGCs whereas the peak density relies on a *subset* of labelled RGCs. Quantification of the developmental dynamics of map formation reveals restricted periods of rapid change.

The two-stage refinement in wild type with an initial slow refinement until P4 and the rapid refinement seen from P4 (Figure 2B) may simply reflect the spatial profile of correlated activity in the retina. In the normal retina this asymptotes for separations greater than 600 µm (Stafford et al., 2009). Our anatomical data indicates that many ganglion cells converging on a single collicular locus at P0 will have a much greater separation in retinal label. Consequently, activity-based segregation will not be discriminatory for those ganglion cells. Indirect support for this argument comes from our modelling studies of wild-type and nAChR-β2^−/−^ mice. In the latter, correlated activity asymptotes at a much greater retinal separation (Stafford et al., 2009; Figure 4A) and the modelling predicts a *more rapid* refinement: more ganglion cells fall within the spatiotemporal correlation window.

When the developmental dynamics were studied further in the nAChR-β2^−/−^ mouse, in which altered patterns of retinal activity in the first postnatal week results in a more diffuse projection in mature mice (McLaughlin et al., 2003), we find that, surprisingly, both measures of map refinement (convergence and segregation) remain largely constant in the first 4 postnatal days and the projection is only subsequently refined (Figure 6B). Although this could indicate that the waves are crucial for the early postnatal refinement, the wild type also exhibits a slow refinement in the same period. The rapid refinement seen in the wild type from P4 onwards is, however, not replicated in the nAChR-β2^−/−^ animals where there is a slower refinement, reminiscent of wild-type refinement prior to P4. The difference is clearly seen in the 10 to 14-fold increase in coverage between wild type and nAChR-β2^−/−^ from P6 to P8 (Figure 6D). For the mutant, the map both remains largely refined by P4 and the strength of correlated activity across the retina, although more extensive spatially, is weaker (Figure 4a; Stafford et al., 2009).

In the formation of retinotopic maps both experimental and theoretical work implicate guidance molecules and patterned neuronal activity. A common view is that molecular cues *define* and activity cues *refine* mappings (McLaughlin and O’Leary, 2005). Important insights have come from studies of the retinocollicular projection in transgenic mice, in which cues have been modified either in isolation or in combination (Cang and Feldheim, 2013). Generally, these have generated descriptions of endpoint mappings. The developmental dynamics of map formation remain under-explored experimentally and computationally.

Substituting nAChR-β2^−/−^ activity patterns for wild-type patterns (Stafford et al., 2009) into the computational model failed to capture the biology: refinement was initially *faster* since the wider spatiotemporal correlation window in nAChR-β2^−/−^ provides positional information at larger distances. Patterned activity may be required to read-out molecular guidance cues (Nicol et al., 2007). By reducing the relative strength of the molecular parameter (15% of normal) we could mimic the delayed refinement observed in vivo. A weaker readout of the gradients in the model means that a pair of RGCs at a given separation is now differentiated by a smaller difference in energy levels than they were before. The probability bias of picking the target location, instead of spurious neighbouring locations is smaller (since it depends on the difference in chemical energy which was downscaled), and the system needs more trials to find the right configuration. Another consequence from the change in the read-out is that the final map is less precise.

One assumption of our current model is that we assume that time in the model can be mapped, at least monotonically, onto developmental time (Figure 4). As originally formulated (Koulakov and Tsigankov, 2004), this class of energy-based model makes no such claims, and so we acknowledge that a future model may require a closer link of developmental time between model and experiment. Another limitation of the model is that it is isotropic in the SC with respect to the ML and AP axes, and so it cannot capture the different dynamics of map organisation observed in mice along those two axes. Finally, we have restricted to ourselves to modelling retinal activity at around P9 when the corresponding retinal waves were recorded (Stafford et al., 2009). This ignores developmental changes in spontaneous activity during the first few weeks of life (Maccione et al., 2014).

By focusing on the dynamics of retinocollicular ordering in experimentation and modelling, we have shown that changes in patterned spontaneous activity have a much more dramatic effect in early development than expected. In this paper, we have investigated changes in just one parameter: spontaneous activity in the first post-natal week. The wild-type data could be used to ask how alterations in other cues, or combination of cues, alter map dynamics and how well computational models can explain these data.

## Materials and Methods

### Animals

Wild-type animals are C57BL6/J (Charles River, UK). nAChR-β2^−/−^ mice are on the C57BL6/J background and carry a null-mutation for the CHRNB2 gene (Picciotto et al., 1995). Genotyping was done using primers as described (Bansal et al., 2000). Neonatal animals were bred in-house from above strains on 12-hour light/dark cycle. Procedures were performed at multiple ages from postnatal day P0 to P22, with P0 being the day of birth, and adult (8-52 weeks) with a variation of ±8 hours. All procedures were performed in accordance with the European Communities Council Directive of 24 November 1986 (86/609/EEC) under the Animals (Scientific Procedures) Act 1986.

### Injections

Mice (P5+) were anaesthetized either with Isoflurane (1.5-2%) or intraperitoneal injection of Ketamine (130mg/kg) and Xylazine (13mg/kg). Neonatal animals (P0-4) were always anaesthetised with intraperitoneal injection of Ketamine (130mg/kg) and Xylazine (13mg/kg). Animals were given a lethal dose of pentobarbital Sodium (300mg/kg) and perfused transcardially with phosphate buffered saline followed by paraformaldehyde (4% w/v) at 24 hours (for neonatal animals) or 48 hours (for adult animals) after procedures.

Injections were performed using pressure and small diameter borosilicate glass pipettes (Plowden & Thompson, UK) pulled to obtain a tip size of 10-20µm. For SC injections, 5-10nl of red or green fluorescent latex microspheres (Lumafluor, USA) (Katz et al., 1984), diluted 1 in 3 with phosphate buffered saline, were injected unilaterally into the superficial SC. All injections were performed using epifluorescence and oriented to be perpendicular to the surface, as described previously (Upton et al., 2007). Paired red and green injections were done at varying locations across the SC and with varying AP and ML separations of between 50μm and 2000μm. The average injection site measured 70±17µm in diameter at the widest point and was 210±55µm deep. In P0 and P2 animals, injections were made through a cranial pinhole. In animals between P4 and P22, a craniotomy was performed and injections were done through the dura. For adult animals and for anterior injections in older neonatal animals (P8-P22), a craniotomy was performed and the cortex was aspirated to reveal the surface of the SC. All injections were placed with reference to the vasculature at the midline and posterior border of the SC and injections were angled to be perpendicular to the surface of the SC.

### Tissue processing

Following perfusion fixation, the brain was removed and imaged whole to obtain the position of the injection sites. Brains were subsequently embedded in 2% gelatine and sectioned into 100µm parasagittal sections. Finally, the SC sections were imaged and the extent and separation of injection sites was measured using ImageJ (NIH, USA). Before removal of the eyes, a nasal orienting cut was made into the iris centrally in the nictitating membrane and parallel to the eyelids. Eyes were then removed and flat-mounted in Fluoromount (Sigma, UK) before imaging and/or tracing. Locations of labelled cells were plotted using a Camera-Lucida setup with custom-written software. The outlines of cells containing red and green fluorescent microspheres were either traced using a 1 in 4 sampling strategy in a 200 µm by 200 µm grid or every grid point was sampled. The retinal outline was traced at 2.5x magnification and the cells were traced at 25x magnification. The presence of double-labelled cells was defined as cells marked in both channels independently with an offset of no more than 5 μm and with similar cell morphologies.

### Retinal analysis

To standardise the retinal flat-mounts, retinal outlines were reconstructed using Retistruct (Sterratt et al., 2013). Briefly, the retinal outline is traced and the cuts and tears resulting from flattening the retina are marked up manually. The Retistruct algorithm matches points along the edge of the cuts to the rim. A grid is triangulated over the surface of the outline, which is then projected onto a curtailed sphere with a rim-latitude reflecting the shape of the retina at a given age. Subsequently, the grid is optimised over several iterations to minimise the distortion energy of the reconstructed retina. See Sterratt et al. (2013) for details of the reconstruction algorithm.

The proportion of retina containing labelled cells is defined as the area enclosed by a polygon connecting all the outlying cells in an area-preserving (Lambert) projection. The distribution of label in the retina was quantified by plotting isodensity contours to the amount of probability mass they exclude, e.g. the 5% isodensity contour excludes cells with the 5% lowest density measure. Isodensity contours were plotted to 5% 25%, 50%, 75% and 95% of the maximum density estimate using kernel density estimates for fully sampled retinae or kernel regression for partially sampled retinae, as described previously (Sterratt et al., 2013).

Label segregation was calculated using a nearest neighbour algorithm that computes the probability of the nearest neighbour to a given cell being the same colour. This is repeated for all cells of both colours and averaged. Complete overlap of label will result in a segregation value of 0.5, whereas non-overlapping label distributions will result in a value of 1. The data was plotted as a function of SC injection site separation and fitted with the Hill function: 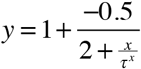 where *y* is segregation and *x* is distance.

Jackknifing (Roff, 2006) was used to estimate the variance of the fits. By calculating a fit while leaving out a data point, and repeating this for all data points, a set of curves is generated. For any *x* the fitted *y* value is the median of all the curves, and the range is specified by the min and the max; the range is visualised as a shaded confidence interval. Non-overlapping confidence intervals for two curves are taken to indicate a significant difference in the two curves.

### Modelling

Wild-type and nAChR-β2^−/−^ retinotopic map formation was simulated using an adapted version (Hjorth et al., 2015) of a previously published model (Triplett et al., 2011). The model tracks the addition and removal of synapses between RGCs and neurons in the SC. Starting from no synapses, a diffuse map initially develops that refines over time. The system has an energy function and favours synapses that reduce total energy. The energy consists of three terms, representing the chemical cues, correlated activity and competition for resources. To reduce the contribution of molecular cues (Figure 8), the energy term *E*_*chem*_ was multiplied by a scale factor less than one. See Triplett et al. (2011) and Hjorth et al. (2015) for details.

In the original model the neurons are placed on a grid; here the neurons are placed randomly using the *d*_*min*_ algorithm, which places an exclusion zone around each neuron, preventing neuronal overlap (Galli-Resta et al., 1997). All parameters and gradients from the original model were used, with the exception of the spatial correlation *b*, and the introduction of the scale factor *μ*.

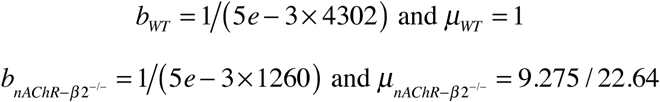

These parameters were changed to account for activity patterns (Stafford et al., 2009) where nAChR-β2^−/−^ activity has a lower correlation than wild type at short distances, but an overall wider correlation window. The values are derived from the assumption that the flattened retinal diameter is 5mm.

To compare results with experimental data virtual retrograde tracer injections were simulated. The virtual injections were matched to the mean size of the experimental injections at 2.8% of SC (data not shown) and the segregation of the marked neurons in the retina was calculated using a similar method as for the experimental data.

To match model iterations to developmental age, the model state was saved at regular intervals. For each saved time point 100 virtual injections were made and a logistic curve fit to the resulting label segregation values. The iteration that best matched the data from a given developmental age was used. Here the summed least square distance between the iterations logistic curve and the experimental data was used to calculate the fit.

## Acknowledgements

This work was supported by a Programme Grant from the Wellcome Trust (G083305) to DJW, IDT & SJE. DL was supported by a Medical Research Council (UK) Studentship. We thank Michael Reber for comments on a draft version of this manuscript.

## Data and code

relating to this paper is available at http://github.com/sje30/b2

